# Subcellular relocalization and nuclear redistribution of the RNA methyltransferases TRMT1 and TRMT1L upon neuronal activation

**DOI:** 10.1101/2020.10.17.343772

**Authors:** Nicky Jonkhout, Sonia Cruciani, Helaine Graziele Santos Vieira, Julia Tran, Huanle Liu, Ganqiang Liu, Russell Pickford, Dominik Kaczorowski, Gloria R. Franco, Franz Vauti, Noelia Camacho, Seyedeh Sedigheh Abedini, Hossein Najmabadi, Lluís Ribas de Pouplana, Daniel Christ, Nicole Schonrock, John S. Mattick, Eva Maria Novoa

**Author notes:** Current address: Brigham and Women’s Hospital, Harvard Medical School, MA, USA. Equal contribution (Nicky Jonkhout, Sonia Cruciani). Correspondence to: John S Mattick and Eva Maria Novoa. Lead Contact: Eva Maria Novoa.

## Abstract

RNA modifications are dynamic chemical entities that regulate RNA fate, and an avenue for environmental response in neuronal function. However, which RNA modifications may be playing a role in neuronal plasticity and environmental responses is largely unknown. Here we characterize the biochemical function and cellular dynamics of two human RNA methyltransferases previously associated with neurological dysfunction, TRMT1 and its homolog, TRMT1-*like* (TRMT1L). Using a combination of next-generation sequencing, LC-MS/MS, patient-derived cell lines and knockout mouse models, we confirm the previously reported dimethylguanosine (m ^2,2^ G) activity of TRMT1 in tRNAs, as well as reveal that TRMT1L, whose activity was unknown, is responsible for methylating a subset of cytosolic tRNA ^Ala^ (AGC) isoacceptors at position 26. Using a cellular *in vitro* model that mimics neuronal activation and long term potentiation, we find that both TRMT1 and TRMT1L change their subcellular localization upon neuronal activation. Specifically, we observe a major subcellular relocalization from mitochondria and other cytoplasmic domains (TRMT1) and nucleoli (TRMT1L) to different small punctate compartments in the nucleus, which are as yet uncharacterized. This phenomenon does not occur upon heat shock, suggesting that the relocalization of TRMT1 and TRMT1L is not a general reaction to stress, but rather a specific response to neuronal activation. Our results suggest that subcellular relocalization of RNA modification enzymes play a role in neuronal plasticity and transmission of information, presumably by addressing new targets.

## INTRODUCTION

Epigenetic processes involving DNA methylation and histone modifications are vital for learning and memory formation (1). RNA modifications, which are much less characterized, add an additional and largely unexplored layer of regulation and avenue for environmental response in cellular function (2–9). Dysregulation of the RNA modification machinery has already been linked to many human diseases (10–13), including several neurological diseases (14–20). A number of studies have pointed to RNA modifications as potential molecular mechanisms underlying the experience-dependent plasticity of the brain (2, 7, 21–23), but a systematic characterization of which RNA modifications may be playing a role in neuronal plasticity and response to environmental signals is lacking.

Cognitive impairment of intellectual disability (ID) is a genetically heterogeneous neurodevelopmental disorder, estimated to affect 2-3% of the population (24). Mutations in three RNA methyltransferase-encoding genes have been linked to intellectual disability, suggesting an important role for RNA modifications in the development of cognitive functions. These include the 2’-O-methyltransferase FTSJ1, identified in X-linked non-syndromic intellectual disability in which mental retardation is the sole clinical feature (25, 26), as well as the m ^5^ C methyltransferase NSUN2 and the m ^2,2^ G methyltransferase TRMT1, both of which have been identified as the cause of autosomal-recessive intellectual disability (ARID) (27–30).

TRMT1 is an RNA methyltransferase responsible for the formation of N2,N2-dimethylguanosine (m ^2,2^ G) at position 26 in both cytosolic and mitochondrial tRNAs (31, 32). In vertebrates, TRMT1 has a homolog, *TRMT1-like* (TRMT1L), in which the methyltransferase domain is conserved (33). The enzymatic activity of human TRMT1 has been previously demonstrated (31, 32), but the function and targets of TRMT1L are unknown. Similarly to TRMT1, the TRMT1L gene is involved in cognitive functioning (33). Indeed, knockout of TRMT1L in mice has been shown to cause altered motor coordination and aberrant exploratory behavior (33), suggesting that its activity is also associated with neurological functions.

To dissect the biological role of TRMT1L, we employed a combined strategy of Liquid Chromatography-Mass Spectrometry (LC-MS/MS) and next-generation sequencing (NGS) to map m ^2,2^ G modifications both in long and short RNAs in TRMT1L knock-out and wild type mouse brain samples. As a control, we also analyzed the m ^2,2^ G modifications in TRMT1-deficient and control cell lines. Through this analysis, we confirm the previously reported dimethylase activity of TRMT1 in cytosolic and mitochondrial tRNAs (31, 32), as well as show that TRMT1L specifically dimethylates a subset of tRNA ^Ala^ (AGC) isoacceptors at position 26. Moreover, using a cellular *in vitro* model that mimics neuronal activation and long term potentiation (34), we report that both TRMT1 and TRMT1L change their subcellular localisation upon neuronal activation, but not upon other stress exposures.

## RESULTS

### Evolutionary analysis of the human m ^2,2^ G modification machinery: TRMT1 and TRMT1L

Human TRMT1 and TRMT1L only share 20% similarity in their DNA sequence (32); however, their domain structure is relatively similar, including a conserved homologous methyltransferase domain (**Figure 1A**). Whilst the dimethylguanosine transferase activity of TRMT1 has been previously reported (31, 32), the activity of TRMT1L is unknown.

**Figure 1.**
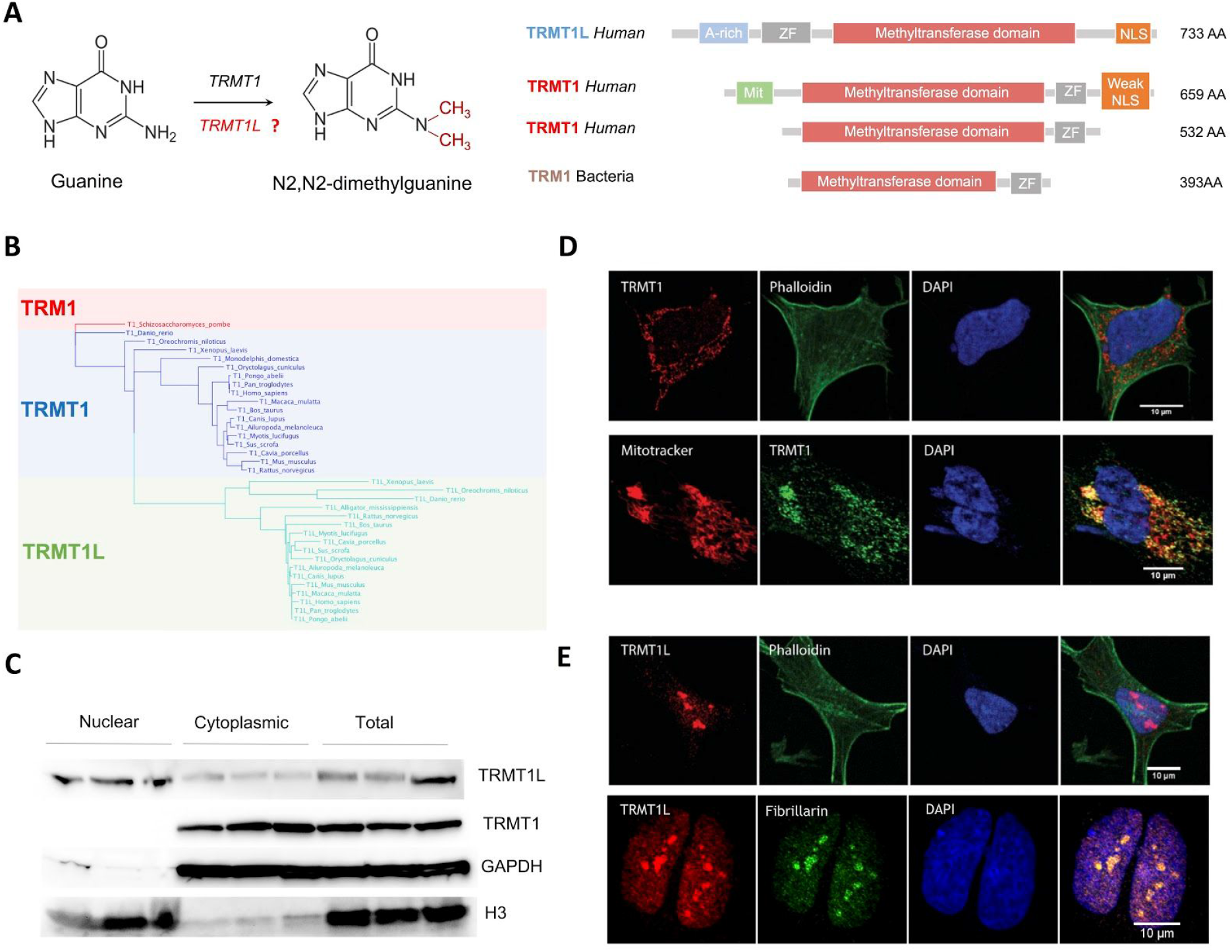
Characterization of the m ^2,2^ G modification machinery. (A) Chemical structure and dimethylation reaction catalyzed by TRMT1 and potentially by TRMT1L is shown in the left panel. In the right panel, examples of the domains found in proteins containing TRM1 methyltransferase domain, such as TRMT1L, TRMT1 and TRM1, are depicted (B) Phylogenetic tree of proteins with homology to the TRM1 methyltransferase domain, showing that TRMT1L proteins are paralogs of TRMT1. (C) Immunoblotting of TRMT1L and TRMT1 in human SH-SY5Y cells, which were fractionated into nuclear and cytoplasmic content. H3 was used as a nuclear marker, whereas GAPDH was used as cytoplasmic marker. Biological triplicates are shown. (D and E) Immunofluorescence of TRMT1 and TRMT1L in SH-SY5Y cells. Nucleus has been marked with DAPI, cytoplasm has been labeled using anti-phalloidin antibody, mitotracker has been used to label the mitochondria, and anti-fibrillarin has been used to label the nucleolus. See Figure S1 for immunofluorescence results in HeLa.

To identify the full set of enzymes potentially capable of placing m ^2,2^ G in the human transcriptome, we performed an HMM-based search of the m ^2,2^ G methyltransferase domain (Pfam domain annotation: TRM), finding that only TRMT1 and TRMT1L showed statistically significant matches in human. We found that the duplication of TRMT1 occurred at the base of vertebrates, originating TRMT1L (**Figure 1B**), which was retained in all vertebrate species analyzed. From our analyses, we find that the TRM m ^2,2^ G methyltransferase domain is found in all eukaryal and archaeal species analyzed, as previously reported (35), but also in a limited set of bacterial phylums (Aquificales and Cyanobacteria), likely acquired via horizontal gene transfer (**Figure S1**).

### TRMT1 and TRMT1L have different subcellular localizations

TRMT1 and TRMT1L are predicted to reside in different organelles based on their signal peptides. Specifically, TRMT1 is predicted to have two major isoforms, one of them containing a mitochondrial-targeting peptide signal, and thus is likely driving m ^2,2^ G methylation of mitochondrial RNAs, whereas the other TRMT1 isoform is predicted to remain in the cytoplasm (**Figure 1A**). By contrast, TRMT1L contains a predicted NLS signal, and is therefore expected to reside in the nucleus.

To confirm the subcellular localization predictions, both Western Blot and immunofluorescence assays were performed. We observed that TRMT1 was located in punctate domains in the cytoplasm and in the mitochondria of neuroblastoma-derived SH-SY5Y cells (**Figures 1C,D**), whereas HeLa cells showed also nuclear localization in addition to cytoplasmic and mitochondrial localization (**Figure S2**). The observed localizations are in agreement with the previously described activity of TRMT1 on both cytosolic and mitochondrial tRNAs (31, 32). By contrast, TRMT1L in SH-SY5Y cells was mainly localized in the nucleus (**Figures 1C,E**), and more specifically, co-localizing with nucleoli (**Figure 1E**).

Therefore, the distinct subcellular locations of these two enzymes suggest that TRMT1 and TRMT1L are addressing different RNA targets.

### m ^2,2^ G is found in tRNAs, but also in higher RNA fractions in neuronal cell lines and tissues

Early *in vitro* studies in human cells showed that TRMT1 is responsible for placing m ^2,2^ G in numerous tRNAs at position 26 (32). More recently, using CRISPR gene knockout systems, it was shown that TRMT1 deletion leads to a loss of m ^2,2^ G in cytoplasmic and mitochondrial tRNAs (31), confirming its activity in a cellular context. However, whether TRMT1 places m ^2,2^ G in substrates other than tRNAs, and what the activity and biological targets of TRMT1L may be, are still open questions.

To decipher whether m ^2,2^ G is found beyond tRNAs, we performed LC-MS/MS analysis of specific RNA pools. Total RNA samples were fractionated into 5 different pools (see *Methods*) corresponding to: (i) total RNA fraction (containing mostly rRNAs), (ii) ribodepleted polyA(+) fraction (containing mostly mRNAs), (iii) ribodepleted non-polyA(+) fraction (containing lncRNAs), (iv) 120-200nt RNAs (containing snRNAs and snoRNAs), and (v) 70-110nt RNAs (containing mostly tRNAs) (**Figure 2A**). For each RNA pool, we quantified the m ^2,2^ G levels using LC-MS/MS, in biological triplicates, and normalized its abundance to the abundance of unmodified bases in each RNA pool. As expected, we were able to detect m ^2,2^ G in the 70-110nt RNA pool, which includes mature tRNAs. In addition, we also detected significant levels of m ^2,2^ G in the 120-200nt RNA fraction (**Figure 2B**), suggesting that m ^2,2^ G is not only found in mature tRNA molecules.

**Figure 2.**
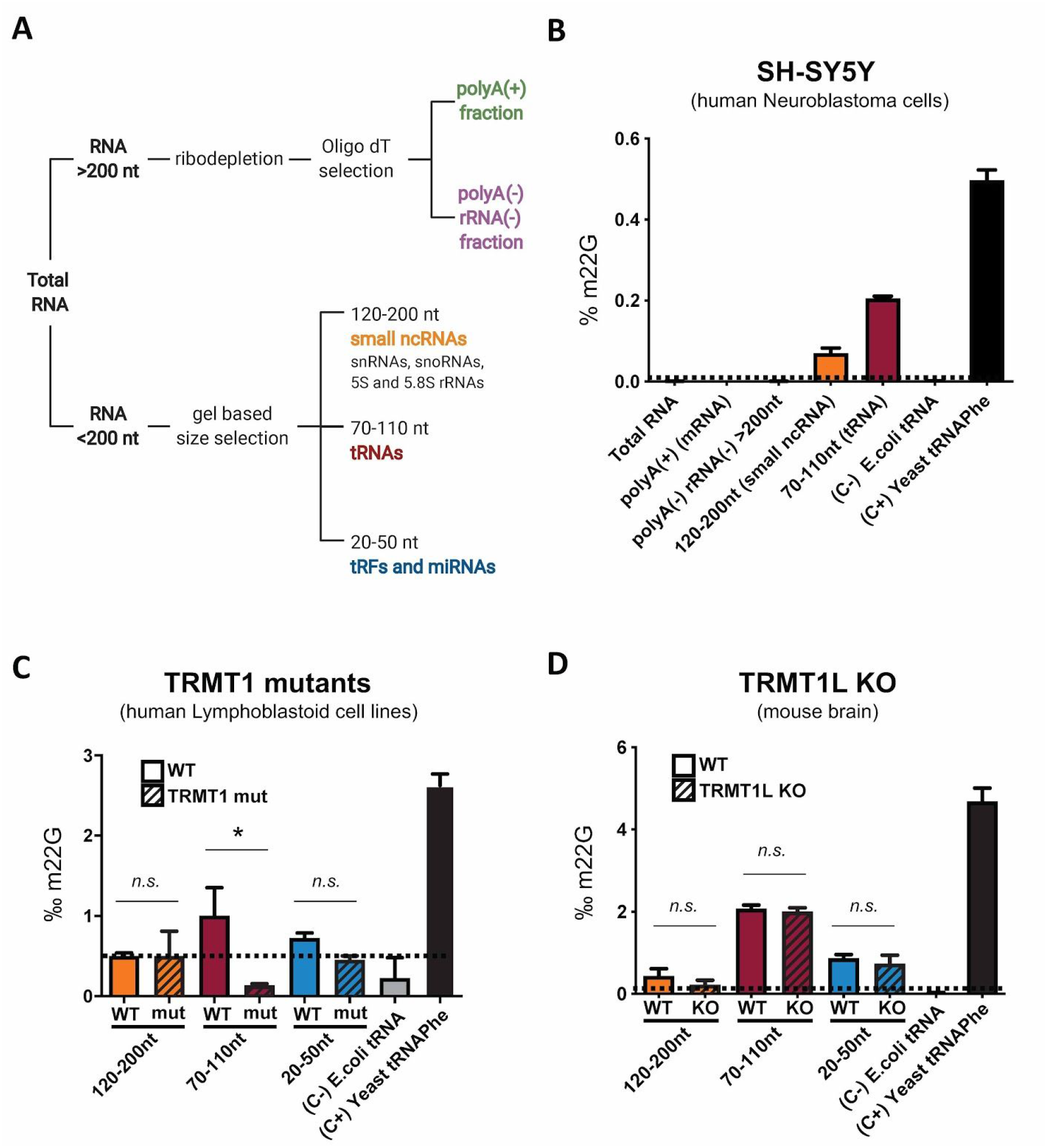
Quantification of m ^2,2^ G in different RNA pools using LC-MS/MS. (**A**) RNA fractionation strategy used for LC-MS/MS samples. (**B**) LC-MS/MS m ^2,2^ G quantification in the 5 different RNA pools of neuron-derived SH-SY5Y cells. Standard deviation denotes technical triplicates. (**C**) Relative proportion of m ^2,2^ G in 3 size selected RNA pools in human TRMT1 mutant patient-derived lymphoblastoid cells, relative to human TRMT1 wild-type patient-derived lymphoblastoid cells. Standard deviation denotes biological triplicates. (**D**) Relative proportion of m ^2,2^ G in 3 size selected RNA pools in TRMT1L KO mice brain samples, relative to WT mice brain samples. Standard deviation denotes biological triplicates. *E. coli* total tRNA was used as negative control (C-), as it does not contain m ^2,2^ G modifications, and *S. cerevisiae* tRNA ^Phe^ was used as positive control (C+), as it is known to contain m ^2,2^ G in position 26.

### LC-MS/MS identifies tRNAs as TRMT1 targets, but does not provide hints on TRMT1L targets

To identify the biological targets of TRMT1 or TRMT1L, we analyzed the m ^2,2^ G levels in TRMT1L knockout and TRMT1-deficient samples, compared to wild-type, using LC-MS/MS (**Figure 2C,D**, see also *Methods*). The data show that TRMT1 is largely responsible for m ^2,2^ G modification in tRNAs, in agreement with previous observations (31, 32). By contrast, TRMT1L knockout did not significantly alter the global m ^2,2^ G modification levels of any of the 3 RNA fractions examined (**Figure 2D**).

We then wondered whether TRMT1L might in fact be responsible for placing an RNA modification different to m ^2,2^ G, which could explain why m ^2,2^ G levels do not significantly vary between TRMT1L knockout and wild type samples. Using LC-MS/MS, we quantified the RNA modification levels of 27 distinct RNA modifications across 7 RNA pools of distinct RNA sizes (**Figure S3**). We found that the depletion of TRMT1L led to modest decrease in the m ^2,2^ G levels in the 120-200nt RNA fraction as well as in the m ^2^ G levels in the 70-110nt RNA fraction (**Figure S3**), although neither of these were found to be significant.

### Mapping of TRMT1L-dependent modifications in the brain transcriptome

Whilst LC-MS/MS is a sensitive methodology that can accurately quantify RNA modifications, it does not identify which RNAs molecules are in fact modified. Therefore, we examined the RNA modification activity of TRMT1L using next-generation sequencing. Indeed, certain RNA modifications, such as m ^2,2^ G, affect the Watson-Crick base pairing moiety, and can be detected in the form of ‘mismatch signatures, in RNA-seq datasets (36–38). The dimethylation of guanosine at the N2 position eliminates the ability of the N2 to function as a hydrogen bond donor, altering its pairing behavior, and consequently affects the misincorporation rate at that given position when reverse transcribed (39, 40). Similarly, previous studies have shown that m ^1^ G and m ^2^ G modifications can be detected in the form of altered mismatch frequencies (41, 42), although the latter only causes moderate misincorporation defects (43, 44). Overall, both m^2^ G and m ^2,2^ G RNA modifications constitute strong candidates for being detected using the non-random mismatch signature strategy in RNA-seq datasets.

We performed both small and long RNA sequencing of mouse brain samples in both TRMT1L knockout and wild-type mice in biological triplicates (**Figure 3A**), using a modified library preparation protocol to capture small RNAs up to 150 nt (see *Methods*). We found that our strategy was able to capture known m ^2,2^ G modification sites in position 26 of multiple tRNAs, in a highly reproducible manner (**Figure 3B**). We observed that the presence of m ^2,2^ G modifications caused characteristic mismatch signatures at the modified sites, mainly in the form of G-to-T mismatches (**Figure 3C**), in agreement with previous reports. By contrast, we were unable to identify mismatch signatures in the vast majority of m ^2^ G-modified sites in mouse tRNAs. Thus, we concluded that our RNA-seq mismatch analysis would be capturing information related to changes in m ^2,2^ G modifications levels, but not from m^2^ G-modified sites.

**Figure 3.**
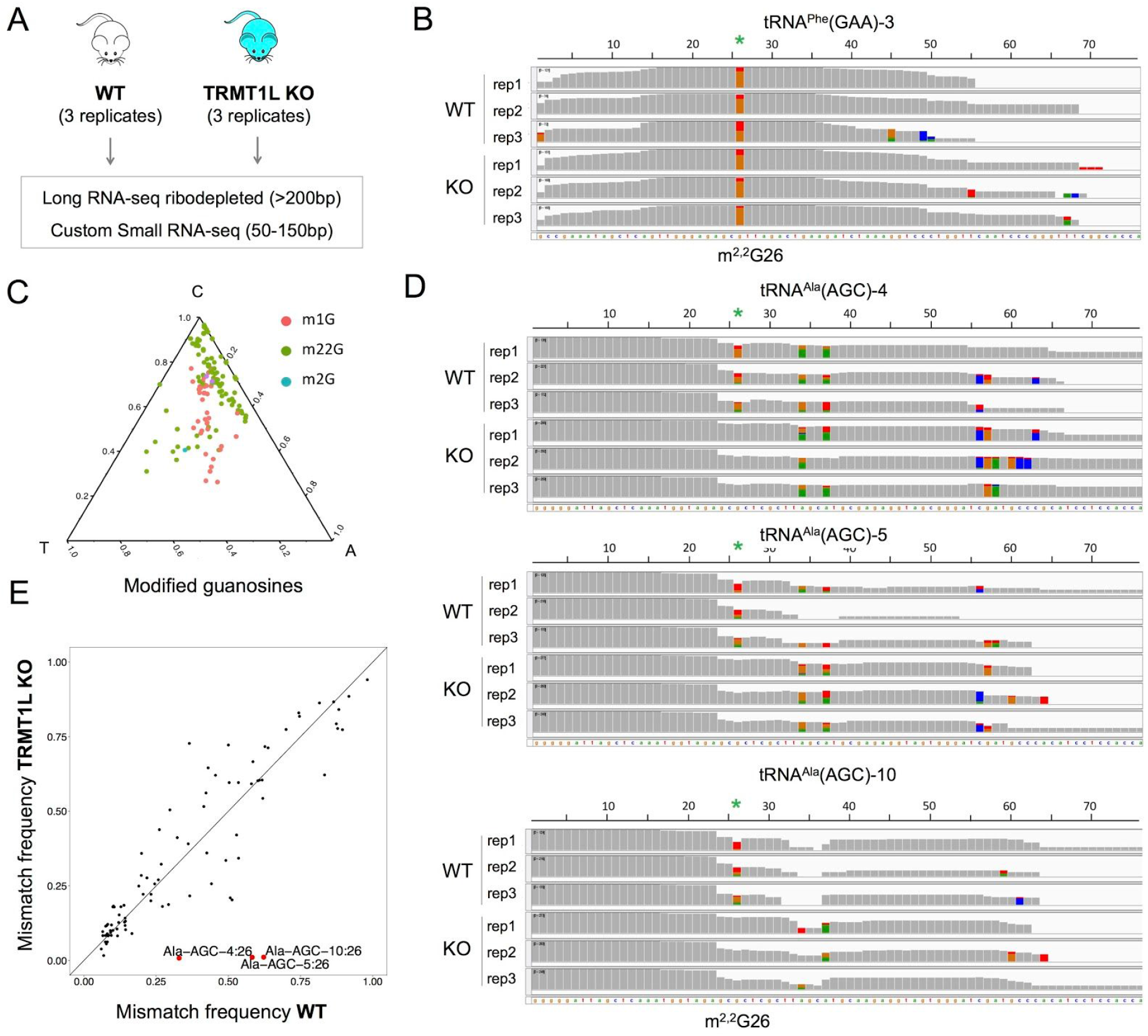
m ^2,2^ G, but not m^2^ G, can be identified via non-random mismatch signatures in RNA-seq datasets. (**A**) Strategy for transcriptome-wide detection of m ^2,2^ G RNA modifications. (**B**) Previously known m ^2,2^ G modification is detected in position 26 of eukaryotic tRNAs. (**C**) Ternary plot of the mismatch signatures of m ^2,2^ G, m ^1^ G and m^2^ G in known tRNA-modified positions. (**D**) IGV tracks of both WT and TRMT1L KO centered on several tRNA ^Ala^ (AGC) gene clusters, showing that the mismatch signatures observed in position 26 consistently disappear in the knockout strain, in all 3 biological replicates. Positions with mismatch frequency greater than 0.1 are coloured, whereas positions with mismatch frequency lower than 0.1 are shown in grey. The m ^2,2^ G26 sites are depicted with green asterisks. **(E)** Comparison of the mean mismatch frequencies observed in tRNA gene clusters of TRMT1L knockout brain samples, relative to wild type. Only tRNA positions with mismatch frequencies greater than 0.1 in the wild type strains have been included in the analysis. The 3 tRNA ^Ala^ (AGC) gene clusters that show differential mismatch frequency in TRMT1L knockout samples are highlighted in red. Mismatches seen in positions 34 and 37 of tRNA ^Ala^ (AGC) correspond to mismatch signatures caused by the presence of inosine (I34) and 1-methylinosine (m1I37), respectively.

We then proceeded with differential mismatch frequency analysis, first focusing on the subset of reads mapping to mature tRNAs (see *Methods*) Our analysis identified 3 tRNA ^Ala^ (AGC) gene clusters whose mismatch error at position 26 completely disappeared in all 3 replicates, suggesting that m ^2,2^ G modifications present in these tRNAs were exclusively placed by TRMT1L (**Figure 3D,E**). We then examined the mismatch signature in reads mapping to other RNA pools, including small non-coding RNAs, pre-tRNAs, rRNAs, lincRNAs and mRNAs; however, we did not identify any significant site whose mismatch frequency changed upon TRMT1L knockout (**Figure S4**).

### tRNA ^Ala^ (AGC) genes modified by TRMT1L are not exclusively expressed in neuronal tissues

Previous studies have shown that a mutation in a CNS-specific tRNA ^Arg^ (UCU) can lead to neurodegeneration in mice (45). The fact that the tRNA ^Arg^ (UCU) gene is only expressed in neuronal tissues explains why the phenotype of the mutant mice was neurological. Following this line of thought, we reasoned the subset of tRNA ^Ala^ genes that is targeted by TRMT1L might be only expressed in neuronal tissues, thus potentially explaining the neurological phenotype observed in TRMT1L knockout mice.

To test this hypothesis, we re-analyzed publicly available small RNA-seq data from mice brain and liver samples across developmental stages (46). Pairwise comparison of tRNA levels in brain and liver tissues identified two tRNAs that were brain-specific: tRNA ^Arg^ (UCU), as previously reported, and tRNA ^Ala^(UGC) (**Figure S5A**). However, we did not identify any tRNA ^Ala^ (AGC) that was brain-specific. Thus, we conclude that the cognitive impairment observed in TRMT1L knockout mice is not a simple consequence of brain-specific expression of the tRNA genes that are being modified by the enzyme (**Figure S5B**).

### Neuronal activation causes subcellular relocalization of the m ^2,2^ G modification machinery

To explore the roles of TRMT1 and TRMT1L in neuronal activity, we compared SH-SY5Y neuroblastoma cells before and after depolarization with potassium chloride (KCl), a system that has been used to mimic long-term potentiation (LTP) (34). Adding 50mM KCl to the medium for 30 seconds causes a short depolarization burst that leads to changes in the expression of immediate early gene (IEG) levels in neuronal cells (47), which have been used as markers for LTP in neuronal cell lines (48). Stimulation-dependent changes in gene expression of the IEGs (C-FOS, ARC, EGR1) were validated by qPCR (**Figure S6**). Under these conditions, we found that TRMT1 relocalizes from the cytoplasm and mitochondria to the nucleus (**Figure 4A**). Likewise, TRMT1L moves from the nucleoli to a similar dotted domain pattern in the nucleus (**Figure 4B,C**). A co-localization experiment showed that, even though TRMT1 and TRMT1L have a similar pattern after activation (**Figure 4D**), they do not colocalize in the nucleus (**Figure 4E**), suggesting that they are being trafficked to different specialized subnuclear domains whose function(s) are yet unknown, indicating the precision and importance of dynamic subcellular localization, the mechanisms of which are also unknown. Moreover, it suggests that these two enzymes may have different RNA targets both before and after activation.

**Figure 4.**
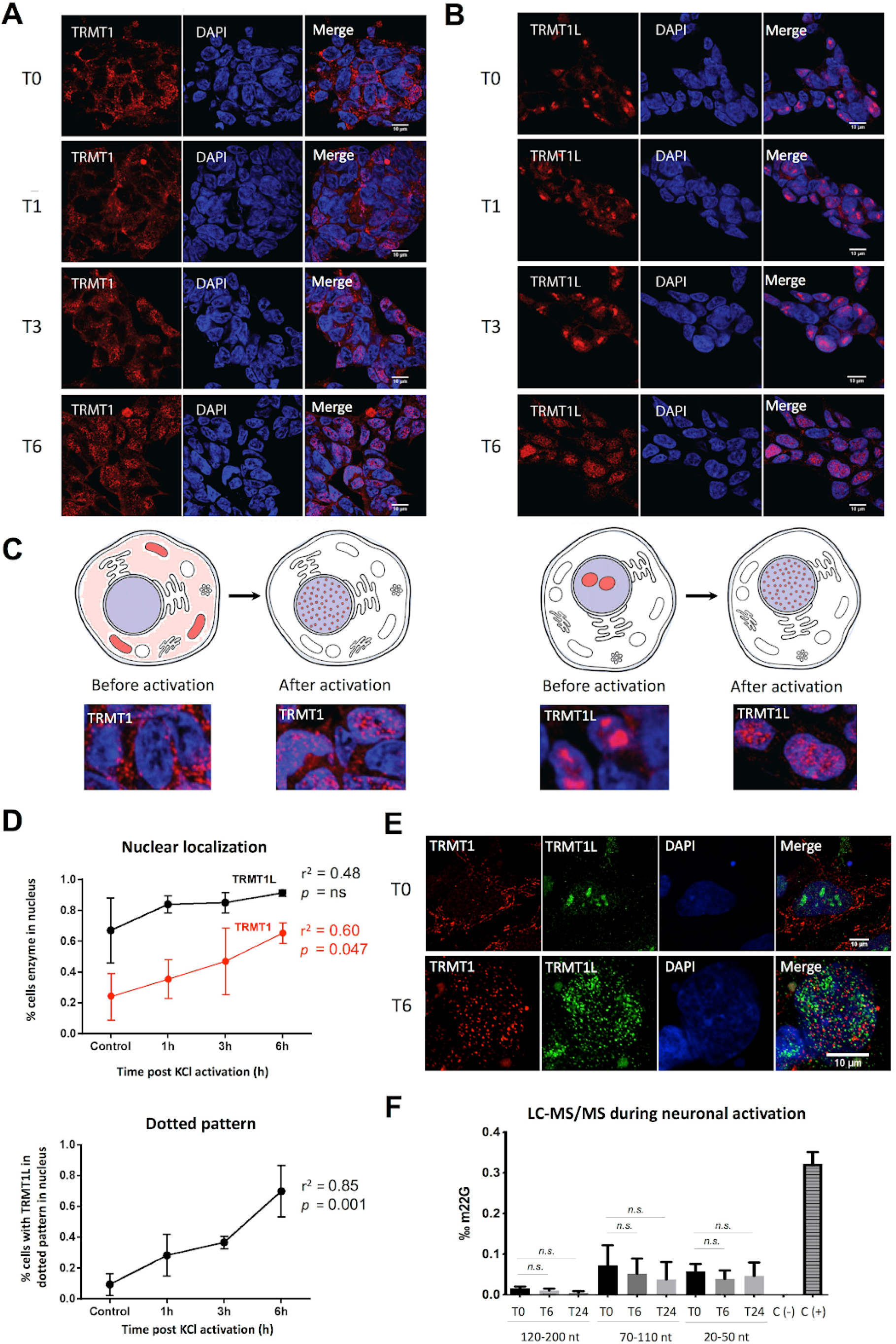
Neuronal activation causes TRMT1 and TRMTL1 subcellular re-localization and affects m ^2,2^ G cellular levels. (**A,B**) Immunofluorescence experiments of SH-SY5Y cells upon neuronal activation, imaged at 0h post-activation (untreated, T0), 1 hour post-activation (T1), 3 hours post-activation (T3) and 6 hours post-activation (T6). SH-SY5Y cells were stained with anti-TRMT1 (A) or anti-TRMT1L (B) antibodies. DAPI was used as a nuclear marker. **(C)** Schematic illustration of the relocation patterns that occur upon neuronal activation, for TRMT1 (left) and TRMT1L (right). (**D**) Relative proportion of cells stained in the nucleus (upper panel) upon neuronal activation. In the bottom panel, the proportion of cells with TRMT1L nuclear dotted pattern upon activation is shown. (**E**) Immunofluorescence of SH-SY5Y cells before (T0) and 6 hours after neuronal activation (T6). Upon activation, TRMT1 relocalizes from cytosol and mitochondrial to the nuclei, but does not colocalize with TRMT1L. (**F**) Abundance of m ^2,2^ G RNA modifications in different RNA pools upon neuronal activation, measured at three time points (t=0h, 6h and 24h post-activation). *E. coli* total tRNA was used as negative control (C-), and yeast tRNA ^Phe^ was used as positive control (C+). Error bars represent standard deviations from 3 biological triplicates.

We then asked whether the relocation of TRMT1 to the nucleus upon neuronal activation might be accompanied by a change in the m ^2,2^ G levels of the cells. To this end, we analyzed the m ^2,2^ G levels of each small RNA fraction individually (120-200nt, 70-110nt and 20-50nt) in SH-SY5Y cells during activation (t=0h, 6h and 24h post-activation). Although we observed a slight decrease in m ^2,2^ G levels upon neuronal activation, these differences were not statistically significant (**Figure 4F**).

Previous studies have shown that the activity of m ^6^ A RNA writer and reader enzymes, as well as the RNA modification levels, vary upon exposure of cells to stress conditions, such as heat shock or chemical stress (49–51). Thus, we investigated whether exposure to heat shock might also cause changes in the localization of TRMT1 and TRMT1L. We found that neither TRMT1 nor TRMT1L change their localization patterns upon exposure to heat shock stress (**Figure S7**) during a 24h time course, in contrast to our observations when exposing SH-SY5Y cells to neuronal stimulation conditions (**Figure 4A,B**). Altogether, our results suggest that the relocalization of TRMT1 and TRMT1L is not a general response to cellular stress, but rather a specific phenotype related to neuronal activation.

## DISCUSSION

Base modifications overlay sequence information to expand the lexicon of RNA. They introduce changes into structural, coding or regulatory information, and are likely to play a major role in the molecular events that underlie brain plasticity and cognitive function. In recent years, it has been shown that some modifications are subject to dynamic regulation (49, 52), can be reversed (11, 52–54), and are involved in many biological processes, including cellular differentiation (55), sex determination (56, 57), maternal-to-zygotic mRNA clearance (58) and stress responses (49, 59). However, much less is currently known about the roles of RNA modification in synaptic plasticity and or memory formation (2).

The most common RNA modification type in the mammalian central nervous system is the deamination of adenosine to inosine (A-to-I), termed RNA ‘editing’, which is catalyzed by adenosine deaminases acting on RNA (ADARs). RNA editing is known to occur in a number of neuroreceptor mRNAs but also thousands of other RNAs, likely playing a major role in neuronal plasticity (60). The extent of RNA editing increases during mammalian and especially primate evolution [1], and the loss of the brain-specific enzyme ADAR3 results in memory deficits and increased anxiety in mice [2]. Moreover, ADARs can shuttle between the nucleus and the cytoplasm upon RNA binding (61), and have been shown to accumulate in the nucleus during neuronal development (62).

In addition to RNA deamination, there are over 100 different RNA base modifications that are formed by addition of chemical moieties (63). Several works have provided genetic evidence that some of these RNA modifications, including ribose, cytosine and guanine methylation, play a role in cognitive function (25–30). However, with the exception of m ^6^ A modifications (21, 64–66), the dynamics of RNA modification in neuronal biology have yet to be studied, largely due to the difficulty of identifying RNA modifications, a problem that is beginning to be solved by the direct RNA sequencing using nanopore technologies (67–70).

In this study, we focused on two RNA methyltransferases that have been previously associated with cognitive dysfunctions: TRMT1, the loss of which causes intellectual disability in humans (27), and its ortholog TRMT1L, the loss of which causes behavioural changes and motor defects in mice (33). We confirmed that TRMT1 is responsible for placing m ^2,2^ G in tRNA molecules, and revealed that TRMT1L is responsible for placing m ^2,2^ G in a subset of tRNA ^Ala^ (AGC) genes. Perhaps not coincidentally, a number of neurological defects result from the loss of tRNAs and related enzymes, whose expression is not necessarily brain-specific (10, 13, 71, 72) and whose effects may be mediated through products of tRNA processing (73).

Most importantly, we show that TRMT1 and TRMT1L are localized in different subcompartments in the cell and that their localization, and presumably their targets, are altered upon neuronal activation. To the best of our knowledge this is the first time that the localization of RNA modification enzymes has been shown to change in response to neuronal activation. The subcellular and subnuclear compartments between which they shuttle (with the exception of the nucleolus) are as yet largely uncharacterized, indicating another unexplored level of neuronal organization, embedded in domains that are likely scaffolded and phase-separated by other RNAs (74, 75). This level of complexity is unprecedented and opens new perspectives in the complex path to the understanding of brain development and neuronal function.

## MATERIALS AND METHODS

### Cell culture and passaging

Human SH-SY5Y neuroblastoma cell lines were cultured in Gibco Life Technologies Dulbecco’s modified Eagle medium (DMEM), supplemented with 10% (v/v) fetal bovine serum (FBS), and 1% Penicillin Streptomycin (Life Technologies, #15070-063). Cells were incubated at 37°C in a humidified atmosphere containing 5% CO_2_. Cells were grown until they reached 80% confluency and washed once with PBS before splitting. For a 150cm2 flask, cells were incubated with 5ml trypsin (Sigma), for 2min at 37°C for detachment. Trypsin was neutralized by adding 15ml of DMEM with 10% FBS. From this mix, 5ml was added to 30ml of DMEM with 10% FBS in a new flask of 150cm^2^, bringing the cells to a confluence of 20%. Cells were discarded when they reached passage number 12.

### Patient-derived TRMT1 mutant and control lymphoblastoid cell lines

B-lymphoid cell lines (LCLs) were established from healthy and affected individuals (**Figure S8**) by *in vitro* infection of peripheral blood mononuclear B cells with Epstein Barr Virus (EBV) based on standard protocols (76). The mutations in the TRMT1 gene (NM_017722) in the patients used to derive the LCLs, as well as the clinical phenotypes of these patients have been previously described (77). Healthy individuals used in this work contained a single copy of the mutated TRMT1 gene and did not show intellectual disability or apparent clinical phenotypes (77), whereas affected individuals contained both TRMT1 gene copies with the 1332_1333 deletion in the coding sequence of TRMT1, and showed intellectual disability (ID) as well as other clinical phenotypes (**Table S1**). All LCL cell lines used in this study were taken from a family with a pY445fs mutation in the TRMT1 coding region (c.1332_1333delGT; p.Y445fs). To generate LCLs, 10ml of peripheral blood was collected from each patient in a sterile preservative-free heparin tube. Mononuclear cells were isolated from whole blood by density Ficoll-Hypaque gradient centrifugation, washed with medium RPMI three times. Pellets were resuspended in transformation-medium (B95-8 virus suspension, RPMI 1640, 20% FBS, 0.2 μg/ml Cyclosporin A). After obtaining appropriate growth, cells were transferred in a 25cm ^3^ filtrated flask, and cultured in RPMI 1640 (Gibco) supplemented with 1% penicillin-streptomycin (Gibco), 15% fetal bovine serum (FBS) (Gibco), 1 mM L-glutamine, 10 mM HEPES and 1% Gibco™ AmnioMAX™ in a 5% CO2, humidified, 37°C incubator up to 3-4 weeks. The culture media were changed every 3-4 days when cells showed extensive growth. Total cellular RNA was isolated from LCLs using VIOGENE miTotal RNA extraction miniprep kit (Cat #VTR1002), in biological triplicates.

### TRMT1L knockout mice brain RNA samples

Trmtl1-like KO mice have been previously generated and described (33). Briefly, ES cells with a gene trap vector integration in intron 1 of the mouse Trmt1-like gene, were used to generate chimeric mice by morula aggregation with wildtype embryos (E2.5) obtained from superovulated CD1-females (Charles River). Following overnight culture of aggregates, blastocysts were transferred to foster mothers and chimeric offspring were mated to C57BL7/6 (Charles River) mice to generate heterozygous Trmt1-like progeny. Heterozygote mice were continuously backcrossed to the C57BL/6 line for more than 10 generations and characterized (33). Tissue samples for this study were obtained from Trmt1-like ^WT/WT^and Trmt1-like ^GT/GT^ mutant mice at the same age of 96-97 days. Mice were culled by cervical dislocation and brains were immediately dissected and snap-frozen in liquid nitrogen and stored at −80°C. Total RNA was obtained by homogenizing whole brain tissues from wildtypes and Trmt1-like mutants using POLYTRON PT 1200E followed by TriZol extraction.

### Immunocytochemistry

Coverslips were placed in 24-well plates, coated with Poly-L-Lysine for 5min and dried overnight after washing 4 times with water. SH-SY5Y cells were plated at a density of 2.5×10^5^ in 500ul of DMEM with 10% FBS and grown overnight. Cells were washed once at room temperature with PBS and fixed in pre-warmed (37°C) 4% paraformaldehyde/4% sucrose in phosphate-buffered saline (PBS) at room temperature for 15min and then washed 3 times for 5min with PBS. Cells were then permeabilized in 0.1% Triton X-100/0.1% Na-Citrate/PBS for 3min at room temperature, and washed 3 times for 5min in PBS. Cells were blocked in 10% FBS/PBS for 1h at room temperature and incubated with anti-TRMT1 primary antibodies (dilution 1:100, Ab134965) or anti-TRMT1L (dilution 1:200, NBP18337) in blocking solution overnight at 4°C. Cells were then washed 3 times for 5min in PBS, and incubated with secondary antibodies for 90min at room temperature, followed by 4 washes with PBS. Coverslips were dipped in demineralized water, and mounted using immuno-fluore mounting medium (MP Biomedicals). Mitochondria staining was achieved using MitoTracker Red FM (Thermo Fisher), by adding MitoTracker 45min before fixation to the growing medium to a final concentration of 100nM. DAPI was employed at a 1:100 dilution, whereas phalloidin was employed at a 1:500 dilution.

### Confocal imaging and image analysis

Confocal images were acquired using a LSM700 confocal laser-scanning upright microscope (Zeiss) with a 63x oil objective. The zoom was set between 1x and 2x, with a pinhole of 34uM and a speed of 1.58us per pixel. Confocal laser intensity was set to 3 and the gain was adjusted per sample. The dimensions were set to 1024×1024pixels and averaged 4 times. All experiments were repeated 3 times and for each condition. The cell counter plugin of Fiji 1.49 (ImageJ) was used to quantify the movement of TRMT1 and TRMT1L after cell activation with KCl. Statistical analysis was done using GraphPad Prism 6, using linear regression analysis.

### Cellular fractionation

Cell fractionation was performed according to the protocol described by Suzuki *et al* (78), with minor modifications. SH-SY5Y cells were grown as monolayers in 15cm diameter dishes until reaching 80% confluence. Cells were washed twice in ice-cold phosphate buffer saline (PBS) pH 7.4 (130mM NaCl, 2mM KCl, 8mM Na_2_HPO_4_, 1mM KH_2_PO_4_), scraped from culture dishes and collected in 1.5 mL of ice-cold PBS. After 15 sec centrifugation in a top table microcentrifuge (Thermo Scientific, USA), cell pellets were resuspended in 750μL of ice-cold 0.05% Nonidet P-40 (Roche Diagnostics, Germany) in PBS and 200μL of the lysate was removed as “whole cell lysate”. The remaining material was centrifuged for 15 sec and 300μl of the supernatant was removed as the “cytoplasmic fraction”. After removal of the remaining supernatant, the pellet was resuspended in 750μL of ice-cold 0.075% NP40 in PBS, centrifuged for 15 sec and the supernatant was discarded. The pellet was designated as the nuclear fraction. Whole cell lysates and cytoplasmic fractions were quantified by measuring OD 280 nm in the Nanodrop 2000 Spectrophotometer (Thermo Scientific, USA).

### Western Blot

Following subcellular fractionation, SH-SY5Y were washed with PBS, harvested and lysed with 70uL of RIPA buffer Samples were then sonicated for 7sec to fragment the DNA for easier loading. LDS Sample Buffer (Novex #B0007) was added and samples were heated to 70°C for 15min. 20ul were loaded onto a Novex Bolt 4-12% Bis-Tris Plus gel, and separated by electrophoresis for 50min at 200V. Following electrophoresis, samples were transferred to PVDF membranes using wet transfer apparatus (Xcell Surelock, Invitrogen) at 30V for 90min. Membranes were blocked in 5% (w/v) nonfat dry milk in TBS-T for 1h at room temperature. Membranes were incubated with anti-TRMT1 (1:10,000, Ab134965) or anti-TRMT1L (1:2,000, NBP-1 88337) antibodies at 4°C overnight with rotation. After washing with TBS-T 3 times for 5min, membranes were incubated for 90min with appropriate horseradish peroxidase-conjugated secondary antibodies at 37°C. Protein bands were visualized using SuperSignal West Pico chemiluminescent Substrate (Thermo #34080) and imaged using Fusion FX (Vilber). Normalization and intensities were analyzed using FusionCapt Advance FX7. For loading controls values were normalized against GAPDH (dilution 1:50.000, Ab8245) and Histone H3 (dilution 1:50.000, Ab1791).

### RT-qPCR

SH-SY5Y cells plated in 6-well plates were lysed using 750ul of QIAzol Lysis reagent (Qiagen) according to manufacturer’s instructions. Total RNA was extracted using the RNeasy mini kit (Qiagen #74104) according to manufacturer’s instructions. First strand cDNA was synthesized from 1ug RNA, using SuperScript III First-Strand Synthesis System (Invitrogen, Life Technologies), and the synthesized cDNA was stored at −20°C until use. RT-qPCR was performed using the Bio-Rad CFX384 Real-Time PCR Detection System using primers (**Table S2**), using GAPDH and PGKI as internal controls. During the extension phase the fluorescent signals were collected and Ct values of the samples were calculated. Transcription levels of TRMT1 and TRMT1L were normalized to controls by the ΔΔCt method (Livak and Schmittgen 2001). Results from triplicate experiments were grouped and an unpaired t-test was used to compare groups.

### Preparation of samples for Mass Spectrometry: RNA isolation

For extracting different fractions of RNA from SH-SY5Y cell cultures, cells were grown until 80% confluency in a 150cm ^2^ flask. Cells were then washed once with PBS, incubated with 5ml Trizol at room temperature for 5min, and collected using a cell scraper. For each mL of sample, after which 300ul of chloroform were added, vortexed for 15s, incubated 5min at RT, and spun down for 15m at 4°C. The clear top phase was moved to a new tube, and the miRNeasy kit (Qiagen #217004) was used to separate the RNA contained in this phase in two fractions: <200nt and >200nt. Briefly, 1VOL of 70% ethanol was added to the RNA phase and vortexed for 5s. The sample was then transferred to a miRNeasy spin column and spinned down for 30s at max speed. This column collects all RNAs larger than 200nt. 0.65VOL 100% ethanol was added to the flow-through of the first spin column, and vortexed. Mix was then added to a second spin column to capture RNAs smaller than 200nt. Both columns were washed with 700ul RWT, 500ul RPE (2x), and 80% Ethanol. Columns were centrifuged and air-dried before adding 30ul RNase free H_2_O.

### Preparation of samples for Mass Spectrometry: RNA fractionation

The >200nt RNA fraction was ribo-depleted using non-overlapping DNA oligonucleotides that are complementary to rRNA 18s and 28s followed by digestion with Hybridase™ Thermostable RNaseH (Lucigen #H39500), as previously described (79). Briefly, 10ug RNA sample was ribo-depleted by adding 10ul of rRNA oligo library (100uM), and incubated with 1x Hybridisation buffer to a total volume of 20ul. The incubation consisted in a down ramp from 95°C to 45°C with −0.1°C/s. When 45°C was reached, RNaseH (2U/ul), MgCl_2_ (20mM) was added and final volume was brought to 40ul, and incubated at 45°C for 30min. After incubation, 10ul of H_2_O was added to bring the final volume to 50ul. Then, 90ul of RNAClean XP beads with 270ul 100% Isopropanol was added for the purification step. Samples were placed on a magnet for 5min and washed twice with 300ul 85% Ethanol. Samples were then air-dried and RNA was eluted in 30ul RNase free H_2_O. After ribodepletion, Dynabeads Oligo(dT)25 (Invitrogen #61002) were used to separate the mRNA fraction. Dynabeads were resuspended thoroughly and placed on a magnet to aspirate the supernatant, and washed with 100ul of Binding Buffer (20mM Tris-HCl, pH7.5, 1.0M LiCl, 2mM EDTA) twice. After washing, Dynabeads were resuspended in 100ul Binding Buffer. Samples were then adjusted to 75ug in 100ul 10mM Tris-HCl pH 7.5, RNase-free H_2_O. 100ul of Binding buffer was added to the samples, and were then incubated at 65°C for 2min to disrupt secondary structures and immediately placed on ice. Then 100ul of washed beads were added and the mix was placed on a rotator for 5min. After rotation, samples were placed on a magnet for 2min and supernatant containing lncRNA was removed and stored on ice. Beads were resuspended in Washing Buffer B by pipetting carefully. Supernatant was again removed on the magnet and this step was repeated once. mRNA was then eluted from the beads with 20ul 10mM Tris-HCl by heating to 80°C for 2min. The samples were then immediately placed on the magnet and eluted mRNA was transferred to a new tube and stored in −80°C. The supernatant containing the lncRNA was precipitated by adding 0.1x the volume in Ammonium Acetate 7.5M and 3x the volume in 100% Ethanol with 1ul of Pellet paint. Samples were vortexed for 10s and incubated overnight at −20°C. The next day samples were spinned down for 30min at 4°C. Pellet was washed twice with 75% Ethanol and air-dried for 15min before eluting in 30ul RNase-free water.

The <200nt RNA fractions were treated with Turbo DNAse (Invitrogen AM2238) according to manufacturer recommendations. After the treatment, RNA samples were recovered on column using the kit RNA Clean & Concentrator (Zymo Research R1017). Samples were further subdivided into three fractions (120-200nt, 70-110nt, and 20-50nt) by electrophoretic separation and gel excision. Briefly, a 15% TBE-Urea gel (Invitrogen, 1.0mm x 10wells, #EC6885BOX) was pre-run at 100V for 30min with 0.5x TBE (Novex TBE running buffer, #LC6675). Approximately 8 ug of <200nt RNA per sample was mixed to an equal volume of 2x loading dye (NEB #B0363A) and incubated at 70°C for 5min. The gel wells were washed from urea, and 20ul per well was added, leaving an empty well between samples to avoid cross-sample contamination. Electrophoresis was carried out at 100V for 2 hours. The gel was then placed on a shaker for 5min in 0.5x TBE, RNase free H2O with 1:10000 SYBR Gold nucleic acid gel stain (Life tech #S11494), and three bands were excised under a UV-light. RNA was extracted from the gel and purified using the kit ZR small-RNA PAGE Recovery (Zymo Research R1070) following the manufacturer recommendations. Samples were eluted from the column in 10 uL RNAse-free water.

### Digestion of RNA into nucleosides for LC-MS/MS

For RNA nucleoside digestion, a maximum of 1ug of sample was mixed with 250U Benzonase Nuclease (E1014-5KU), 200mU Phosphodiesterase I (P-3243, Sigma), 200U Alkaline phosphatase (P-7923-2KU, Sigma), 20mM Tris-HCl buffer pH 7.9, 100mM NaCl. 20mM, and H_2_O up to 50ul, and incubated for 6 hours at 37°C. After incubation samples were stored in −80°C until the LC-MS/MS experiment. tRNA ^Phe^ from *S. cerevisiae* (Sigma, R4018) was used as positive control (m ^2,2^ G-positive), and total tRNA from *E. coli* (Sigma, R1753) was used as negative control (m ^2,2^ G-negative).

### LC-MS/MS analysis - m22G modifications

The UHPLC-MS/MS system consisted of an Ultra High Performance Liquid Chromatography Accela Pump (Thermo Fisher Scientific, Waltham, USA) and HTC PAL autosampler (CTC Analytics, Zwingen, Switzerland) coupled directly to a TSQ Quantum Access triple quadrupole mass spectrometer (Thermo Fisher Scientific) via an electrospray interface. Liquid chromatography was performed on an Acquity UHPLC HSS T3 Column, 2.1 x 100 mm, 1.8 μm (Waters, Milford, USA). 10 μL of the sample was analysed using gradient elution with aqueous 0.1% formic acid (Solvent A) and 0.1% formic acid in acetonitrile (Solvent B) at a flow of 0.4 mL/min over 7 minutes. The acetonitrile gradient increased linearly from 0% at 1.5 minutes to 100% at 6 minutes and held for 0.5 minutes before returning to 0% at 6.60 and remaining there until 8 minutes. The divert valve was set to waste during the first 0.6 minutes of the run. Analysis was performed in positive ionisation mode using selected reaction monitoring (SRM). Mass spectrometric conditions were optimized for sensitivity using infusions of analyte standards. The sheath gas pressure and the auxiliary gas pressure were 30 and 5 (Thermo Fisher arbitrary units) respectively. The probe temperature was 200C, the capillary temperature was 330C and the tube lens offset was 90V. The mass to charge transitions and collision energies used to detect and quantify the analytes monitored were, for adenosine (A): 268 (136 at 17 V), cytidine (C): 244 (112 at 12 V), guanosine (G): 284 (152 at 14 V), uridine (U): 245 (113 at 10V), and d-(dimethylamino)guanosine (m2,2G): 312 (180 at 15V). 1 alternate SRM transition was also acquired for each analyte as a qualifier to assist if required in the case of interference. Standards used for calibration curves included: adenosine (Sigma-Aldrich, A9251), cytidine (Sigma-Aldrich, C122106), guanosine (Sigma-Aldrich, G6752), uridine (Sigma-Aldrich, U3750), 2-(dimethylamino)guanosine (Santa Cruz, sc-220667). Data processing of chromatograms was performed using the Quanbrowser function of the Xcalibur Software package version 2.0.7 (Thermo Fisher Scientific). Quantification was performed using an external calibration curve over the range 10pg to 200ng for A, C, G, U and 10pg to 40ng for m2,2G. Samples were analysed in duplicate and the mean reported. Calibration curves were run at the start of each batch and then after approximately every 30 samples. The procedure described above applies to results shown in Figure 2 and 4.

### LC-MS/MS analysis - panel of RNA modifications

The ribonucleosides were purified with HyperSep Hypercarb SPE Spin Tips (Thermo Fisher Scientific) prior to LC-MS/MS analysis.Samples were analyzed using an LTQ-Orbitrap XL mass spectrometer (Thermo Fisher Scientific, San Jose, CA, USA) coupled to an EASY-nLC 1000 (Thermo Fisher Scientific (Proxeon), Odense, Denmark). Ribonucleosides were loaded directly onto the analytical column and were separated by reversed-phase chromatography using a 50-cm homemade column with an inner diameter of 75 μm, packed with 4 μm Hydro-RP 80 Å (Phenomenex cat # 04A-4375), as previously described (80). Chromatographic gradients started at 95% buffer A and 5% buffer B with a flow rate of 300 nl/min for 5 minutes and gradually increased to 20% buffer B and 80% buffer A in 40 min. After each analysis, the column was washed for 10min with 20% buffer A and 80% buffer B. Buffer A: 20mM Ammonium Acetate pH 4.5. Buffer B: 95%ACN/5% 20mM Ammonium Acetate pH 4.5. The mass spectrometer was operated in positive ionization mode with nanospray voltage set at 2 kV and source temperature at 200°C. Full MS scans were set at 1 microscans with a resolution of 60,000 and a mass range of m/z 100-700 in the Orbitrap mass analyzer. Fragment ion spectra were produced via collision-induced dissociation (CID) at normalized collision energy of 35% and they were acquired in the ion trap mass analyzer. Isolation window was set to 2.0 m/z and activation time of 10 ms. All data was acquired with Xcalibur software v2.1. Serial dilutions were prepared using commercial pure ribonucleosides (1-2000 pgl, Carbosynth, Toronto Research Chemicals) in order to establish the linear range of quantification and the limit of detection of each compound. A mix of commercial ribonucleosides was injected before and after each batch of samples to assess instrument stability and to be used as external standard to calibrate the retention time of each ribonucleoside. Acquired data were analyzed with the Skyline-daily software (v20.1.1.83) and extracted precursor areas of the ribonucleosides were used for quantification. From the mix of commercial ribonucleosides used, m ^3^ C and m ^5^ C were excluded from the downstream analyses due to their strong overlap in retention time and fragmentation patterns. The procedure above applies to results shown in Figure S3.

### Activation of SH-SY5Y cells via KCl treatment

Cells were plated on sterilized coverslips in 24 well plates for imaging at a density of 3×10^5^cells, or in 6 well plates for RT-qPCR. Cells were activated at 60% confluence by adding 50mM KCl to the medium. After 30 seconds, the medium was replaced with fresh medium. Cells were harvested and fixed at 0, 1, 3 and 6-hour post-activation. SH-SY5Y cell activation was validated using RT-qPCR, which confirmed the upregulation in the expression of immediate-early genes (Hoffman et al. 1993; Okuno 2011; Minatohara et al. 2015).

### Small and long RNA-seq library preparation and sequencing

Total RNA was extracted from TRMT1L KO and wild-type brain samples using TriZol. Both wild-type of TRMT1L KO brain mice were prepared for TruSeq Total RNA Illumina sequencing, in biological triplicates, for both the small (<200nt) and large RNA fractions (>200nt). Briefly, first-strand cDNA was synthesized using SuperScript II Reverse Transcriptase. RNA template was then removed, and second cDNA strand was synthesized using Second Strand Marketing MasterMix from the Illumina kit, cDNA was washed using AMPPure-Xp beads, and 3,ends were adenylated with A-Tailing Mix from the Illumina kit. Adapters containing Illumina barcodes for multiplexing were ligated to the end of the double stranded cDNA, and cDNA was washed and purified using AMPPure beads. DNA fragments containing adaptors on both sides were amplified using 15cycles of PCR. Size and purity of the sample was analyzed on the Agilent 2100 Bioanalyzer, Sequencing was performed using the Illumina HiSeq 2500 platform with 125bp paired-end sequencing in the case o long RNA-seq datasets (>200nt), and with 100bp single-end reads in the case of small RNA-seq datasets (<200nt).

### Bioinformatic analysis of small RNAseq libraries

Small RNA-seq reads were processed with a mapping pipeline adapted from a previous tRNA mapping pipeline (81), with minor changes. The adapted version of the pipeline is publicly available in GitHub (https://github.com/novoalab/tRNAmap_Hoffmann_adapted). Briefly, reads were trimmed with bbduk (from bbmap 36.14) keeping reads with length 15-100 nt. Trimmed reads were then mapped with Segemehl 0.2.0-418 (82) to a modified m38 genome complemented with tRNA genes and pseudogenes masked, and pre-tRNA genes appended as additional chromosomes. Segemehl options of this first mapping were: accuracy=80, differences=3. Reads mapping exclusively to the pre-tRNA reference (not mapping to other genomic sites) were kept for mapping to unique sequences of mature tRNAs (downloaded from gtRNAdb http://gtrnadb.ucsc.edu/genomes/eukaryota/Mmusc10/ on October 2019). This second mapping was performed with Segemehl with accuracy=85 and differences=3. Multimapping reads were kept only if they were showing the same mismatch profile with the different mapping loci (“phased” multimapping handling), as done in the original tRNA mapping pipeline (81). In addition to tRNA mapping, we also analyzed the mismatch signatures of other ncRNAs. For the ncRNA analysis, the trimmed reads were mapped with Segemehl 0.2.0-418 to a custom reference of selected types of ncRNAs (miRNA, scaRNA, snoRNA and snRNA) downloaded from BioMart (GRCm38.p6) with default options, accuracy=95 and differences=0.

### Bioinformatic analysis of long RNAseq libraries

125 bp paired-end long RNASeq reads were processed with Cutadapt v.1.9.1(83) with (adapter sequence: AGATCGGAAGAG, with -m 1 option to exclude empty reads). Reads were aligned with STAR (84) version 2.7.0f with default parameters to mouse genome (GRCm38) with vM21 gencode annotation. Long RNAseq reads were also mapped to mouse canonical rRNA sequences using Bowtie2 (local mode, default settings, -N 1).

### Detection of m ^2,2^ G modifications using mismatch signatures

Modified positions were identified for both small and long RNA mapped reads through the generation of mpileup files using the Samtools mpileup function (85). Mpileup files were further processed with an in-house script (https://github.com/novoalab/mpileup2stats), which generates frequency tables of mismatches, insertions, deletions and RT-drop offs using mpileup files as input. Only positions with a minimum coverage of 10reads/base were considered for downstream analysis. m ^2,2^ G candidate sites were identified as those with a mismatch frequency difference (WT-KO) greater than 25%.

### Ethics statement

Ethical approval for the patient-derived samples from TRMT1 mutant was waived by the Ethics Committee of the University of Social Welfare and Rehabilitation Sciences, Tehran, Iran (IR.USWR.REC.1395.356). Written informed consent form was obtained from legally authorized representatives (the living parents of patients) before the study.

## RESOURCE AVAILABILITY

### Lead contact

Further information and requests for resources and reagents should be directed to and will be fulfilled by the Lead Contact, Eva Maria Novoa.

### Materials Availability

This study did not generate new unique reagents

### Data and Code Availability

Small RNA-seq and long RNA-seq data from TRMT1L knockout and WT brain mouse samples have been deposited in Gene Expression Omnibus (GEO), under the accession codes GSE152429 (small RNA-seq) and GSE156858 (long RNA-seq). The pipeline used to map the reads to tRNA sequences has been made publicly available in GitHub (https://github.com/novoalab/trna_align_hoffmann). The code used to analyze the mismatch frequencies has been made publicly available in GitHub (https://github.com/novoalab/mpileup2stats).

### AUTHOR CONTRIBUTIONS

NJ performed the majority of wet lab experiments described in this work, including immunofluorescence, cell culturing, neuronal activation, RNA fractionation and preparation of samples for Mass Spectrometry and their corresponding data analysis. SC prepared TRMT1L knockout and wildtype samples for Mass Spectrometry in the Orbitrap. SC and EMN performed bioinformatics analysis of the next-generation sequencing libraries. GV and JT contributed with the setup of the gel fractionation experiments and Mass Spectrometry sample preparation. GL helped with the initial stages of bioinformatic data analysis. NC and LRP cultured patient-derived TRMT1 wild-type and mutant lymphoblastoid cell lines and extracted the RNA for Mass Spectrometry analyses. RP performed the Mass Spectrometry sample processing on the Quantum Access. DK prepared the next-generation sequencing libraries. NS, JSM and EMN conceived the project. JSM and EMN supervised the project. NJ, SC and EMN built the figures. NJ, JSM and EMN wrote the paper, with the contribution from all authors.

## ACKNOWLEDGEMENTS

We thank all the members from the Mattick and Novoa labs for their valuable insights and discussion. NJ was supported by a UNSW International PhD fellowship. SC was supported by a fellowship from “la Caixa” Foundation (LCF/BQ/DI19/11730036). This work was supported by NHMRC funds (Project Grant APP1070631 to JSM), funds from the Australian Research Council (DP180103571 to EMN) and funds from the Garvan Young Investigator Award (to NS). This work was partly supported by the Spanish Ministry of Economy, Industry and Competitiveness (MEIC) (PGC2018-098152-A-100 to EMN). Mass spectrometric results were obtained at the Bioanalytical Mass Spectrometry Facility within the Mark Wainwright Analytical Centre of the University of New South Wales, using infrastructure provided by NSW Government co-investment in the National Collaborative Research Infrastructure Scheme (NCRIS) subsidized access to this facility is gratefully acknowledged. The mass spectrometric analyses shown in Figure S3 were performed in the CRG/UPF Proteomics Unit which is part of the of Proteored, PRB3 and is supported by grant PT17/0019, of the PE I+D+i 2013-2016, funded by ISCIII and ERDF. This work is *in memoriam* of Nicky Jonkhout, first author of this work, our colleague and friend. Nicky went on sick leave in June 2019. While he kept struggling back and forth, he persevered until the very end in his intention to return to research and pursue his PhD, with a love for science we could have never had imagined. We would like to thank and remember Nicky for all his passion, effort and the superb skill that he put into this work. Unfortunately he did not live to see this work in its finished form, but his legacy and findings will remain, which we hope will open new avenues in the fields of neurobiology and epitranscriptomics. We will greatly miss him.

## Notes

### Competing Interest Statement

The authors have declared no competing interest.

