## Supplementary Figures for "Subcellular relocalization and nuclear redistribution of the RNA methyltransferases TRMT1 and TRMT1L upon neuronal activation"

[1] Garvan Institute of Medical Research, Darlinghurst, 2010, NSW, Australia.
[2] School of Biotechnology and Biomolecular Sciences, University of New South Wales, Sydney NSW 2052, Australia.
[3] Center for Genomic Regulation (CRG), Barcelona, Spain.
[4] University Pompeu Fabra (UPF), Barcelona, Spain.
[5] Bioanalytical Mass Spectrometry Facility, Mark Wainwright Analytical Centre, University of New South Wales, Sydney, NSW, 2052, Australia.
[6] Departamento de Bioquímica e Imunologia, Universidade Federal de Minas Gerais, Belo Horizonte, Minas Gerais, Brazil
[7] Division of Cellular & Molecular Neurobiology, Zoological Institute, Technische Universität Braunschweig, 38106 Braunschweig, Germany
[8] Institute for Research in Biomedicine, c/ Baldiri Reixac 15-21, 08028 Barcelona, Catalonia, Spain.
[9] Genetics Research Center, University of Social Welfare and Rehabilitation Sciences, Tehran, Iran.
[10] Kariminejad-Najmabadi Pathology & Genetics Center, Tehran, Iran.
[11] Catalan Institution for Research and Advanced Studies, Passeig Lluís Companys 23, 08010 Barcelona, Catalonia, Spain.
[#] Current address: Brigham and Women's Hospital, Harvard Medical School, MA, USA.
[$] Equal contribution

**SUPPLEMENTARY FIGURES**

**Figure S1.** Phylogenetic tree of the TRM1 methyltransferase-related domains across the tree of life, identified using HMM-based homology searches. Proteins have been coloured according to both their predicted methyltransferase function (Trmt1/Trmt1L, Trmt5, PrmA), as well as their domain of life (Eukarya, Bacteria, Archaea). We mainly find homologous $m^{2,2}$G-methyltransferases (green) in eukaryal and archaeal kingdoms, but also in a limited set of bacterial phylums (Aquificales and Cyanobacteria), likely acquired via horizontal gene transfer.

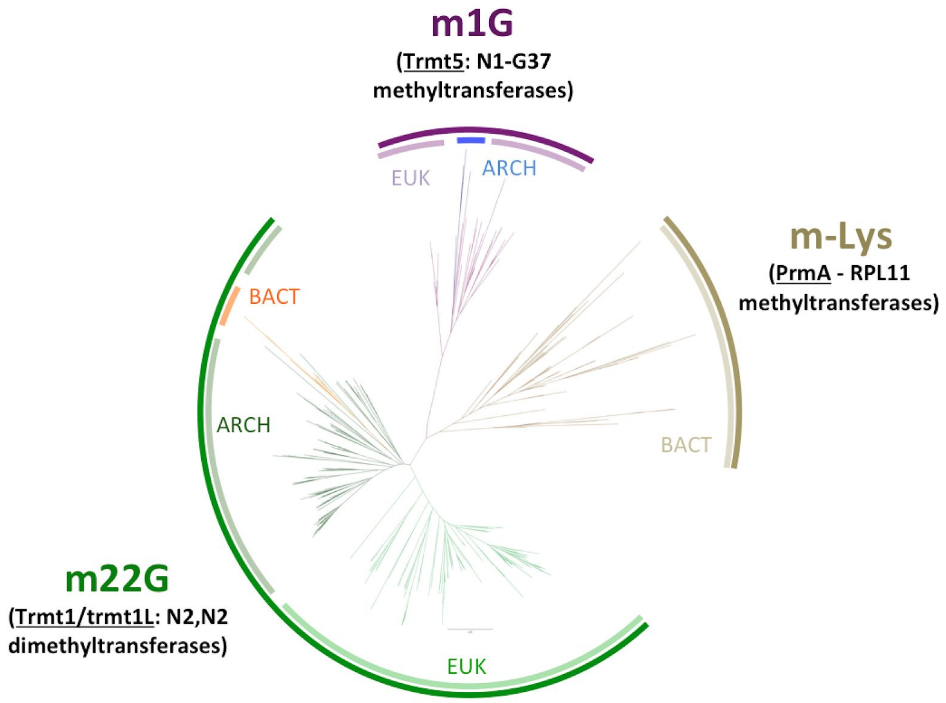

**Figure S2.** Immunofluorescence experiments in HeLa cells show that TRMT1 (green) colocalizes with mitotracker (red). DAPI was used to stain the nuclei (shown in blue).

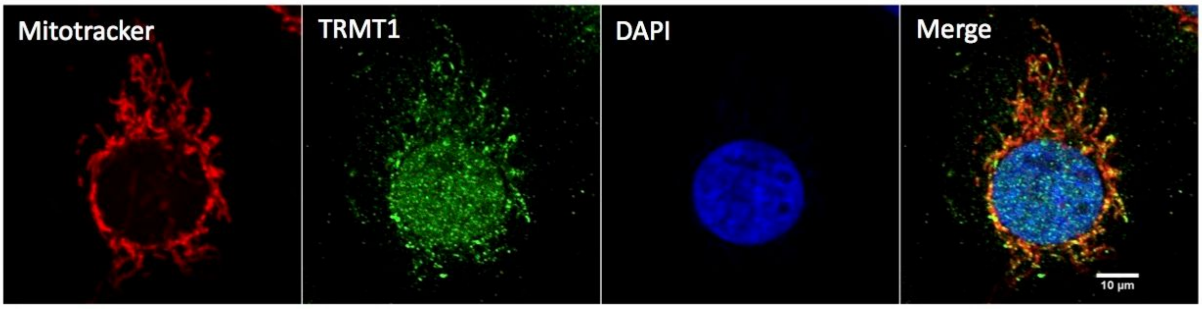


**Figure S3.** RNA modification levels of 27 distinct RNA modifications of TRMT1L knockout and wild type mouse brain samples, across distinct gel-size selected small RNA fractions (<200nt).

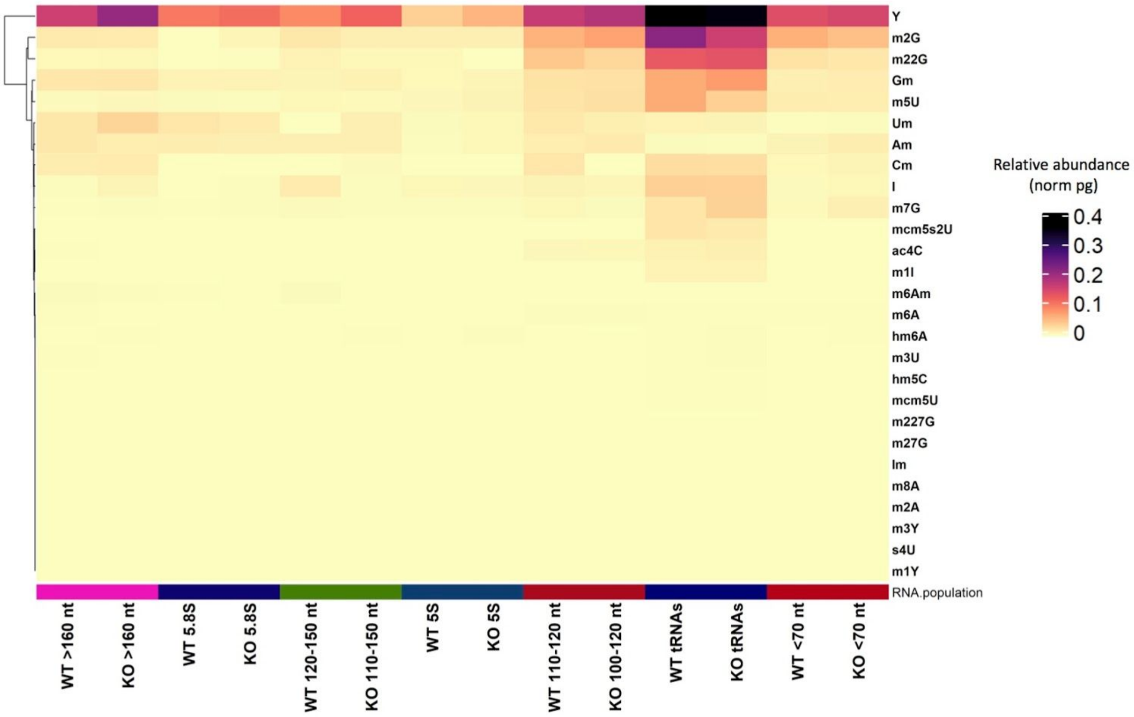

**Figure S4**. Comparative mismatch frequency analysis in TRMT1L knockout and wild type mouse brain samples from small and long RNAseq datasets, in distinct RNA pools.

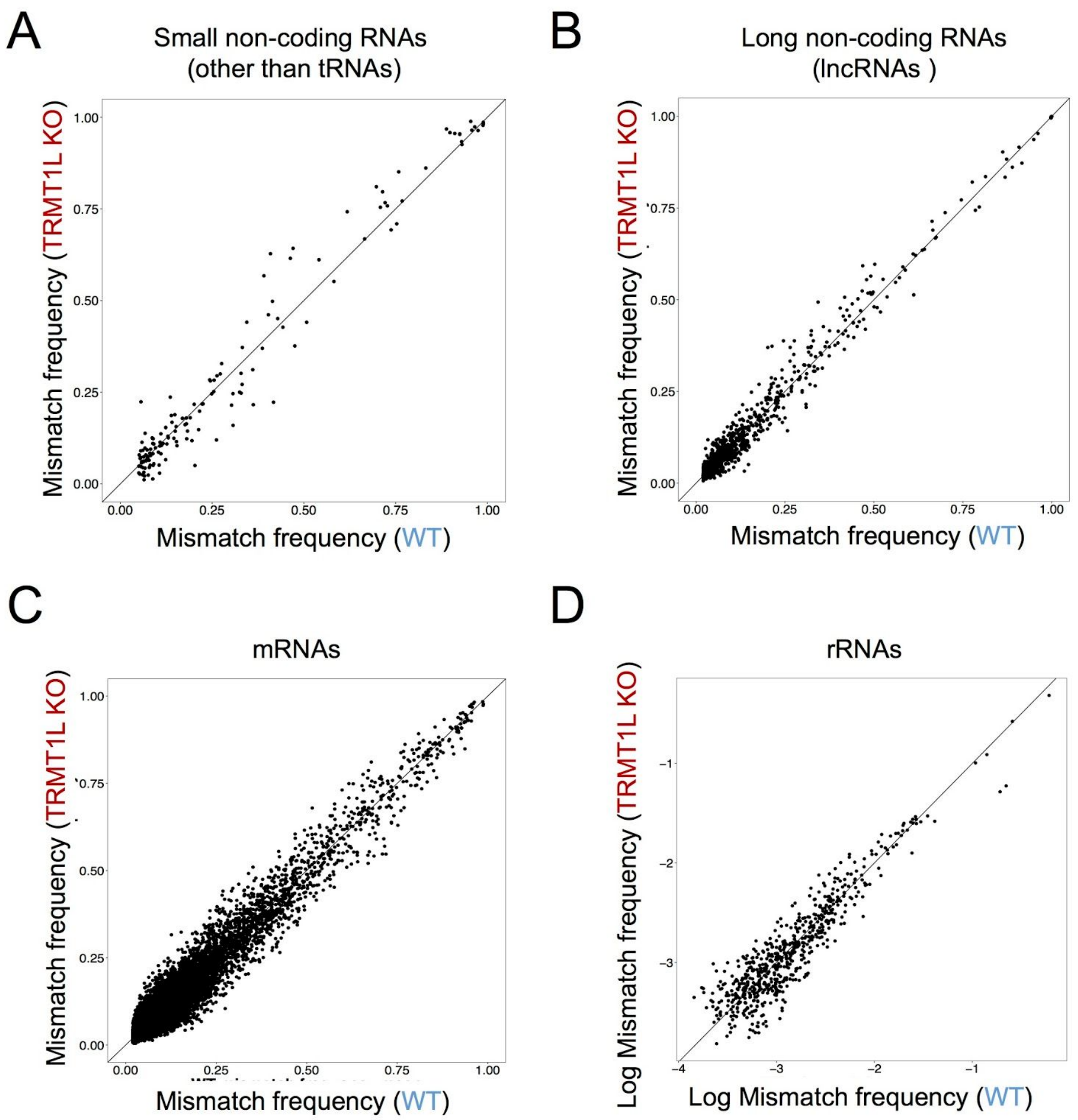

**Figure S5.** Analysis of tissue-specific tRNA expression in brain and liver mice samples from distinct developmental stages. **(A)** Comparison of tRNA counts from brain and liver tissues identifies two tRNA genes tRNA$^{Ala}$(UGC) and tRNA$^{Arg}$(UCU)) that are expressed in the mouse brain but not in the liver. (B) Heatmap depicting the dynamics of tRNAAla (left) and tRNAArg (right) gene expression levels across distinct developmental stages. tRNA genes that show brain-specific expression are boxed in blue.

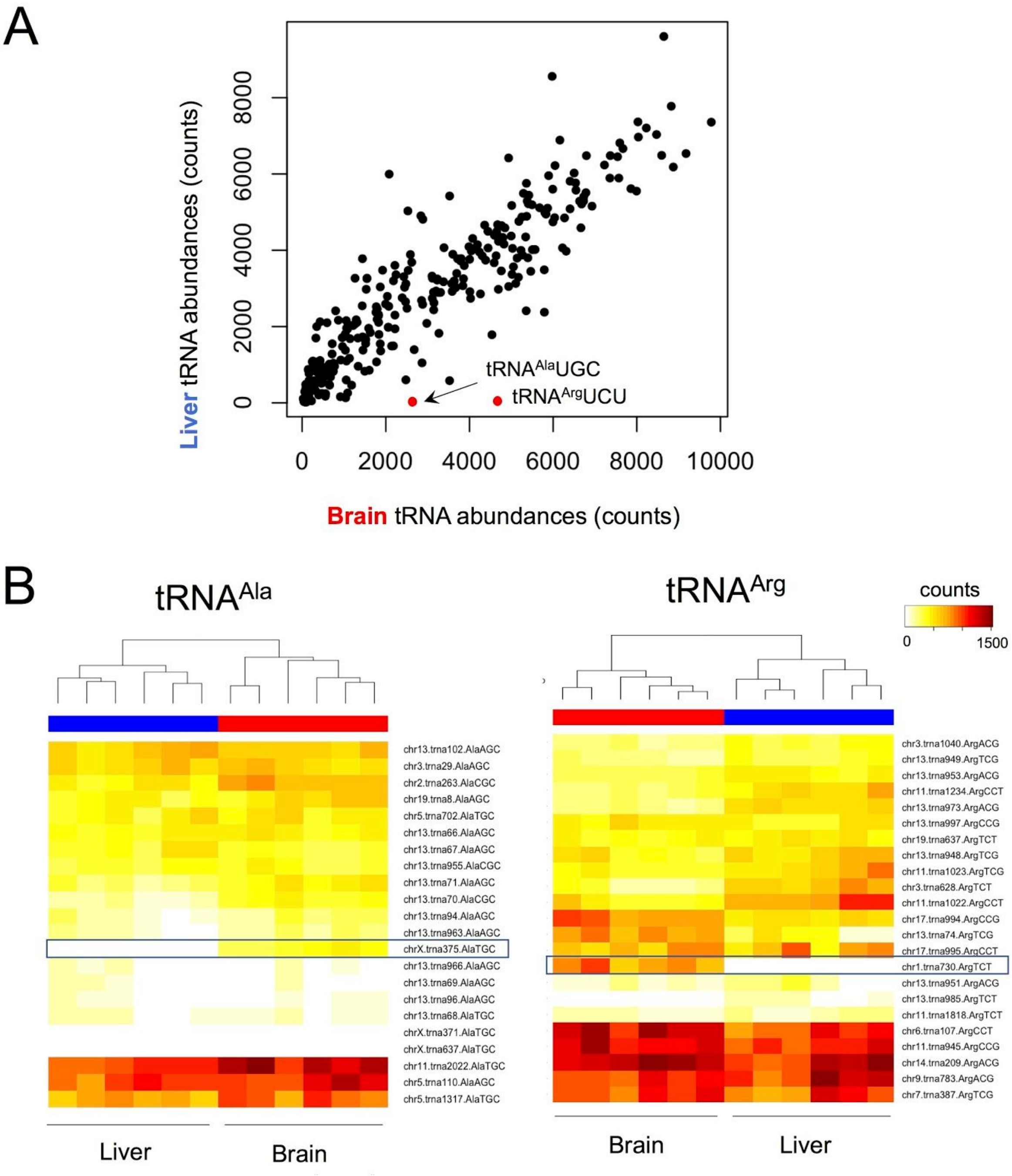

**Figure S6.** RT-qPCR of IEG genes, showing that their expression is increased after neuronal activation, recovering the basal expression levels a few hours post-activation. The expression levels of all genes have been normalized to GAPDH.

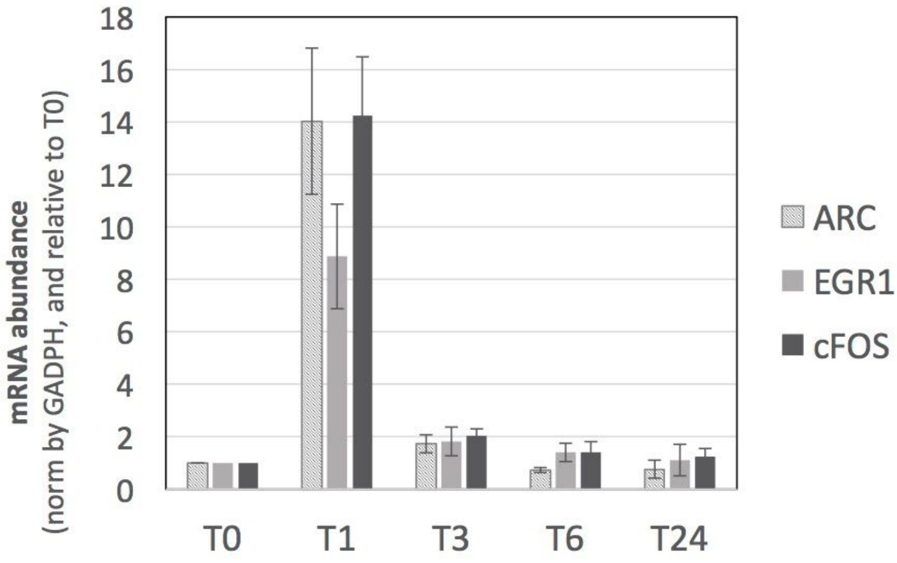

**Figure S7.** Immunofluorescence of SH-SY5Y cells in a time-course experiment upon heat shock exposure, including untreated cells (T0), and heat-shock treated cells at 2h, 6h, and 24h post-exposure to heat shock (t=2h, 6h and 24h). Neither TRMT1 (panel **A**, red) nor TRMT1L (panel **B**, red) relocalize upon exposure to heat shock. Phalloidin has been used to stain the cytoplasm, whereas DAPI has been used to stain the nuclei.

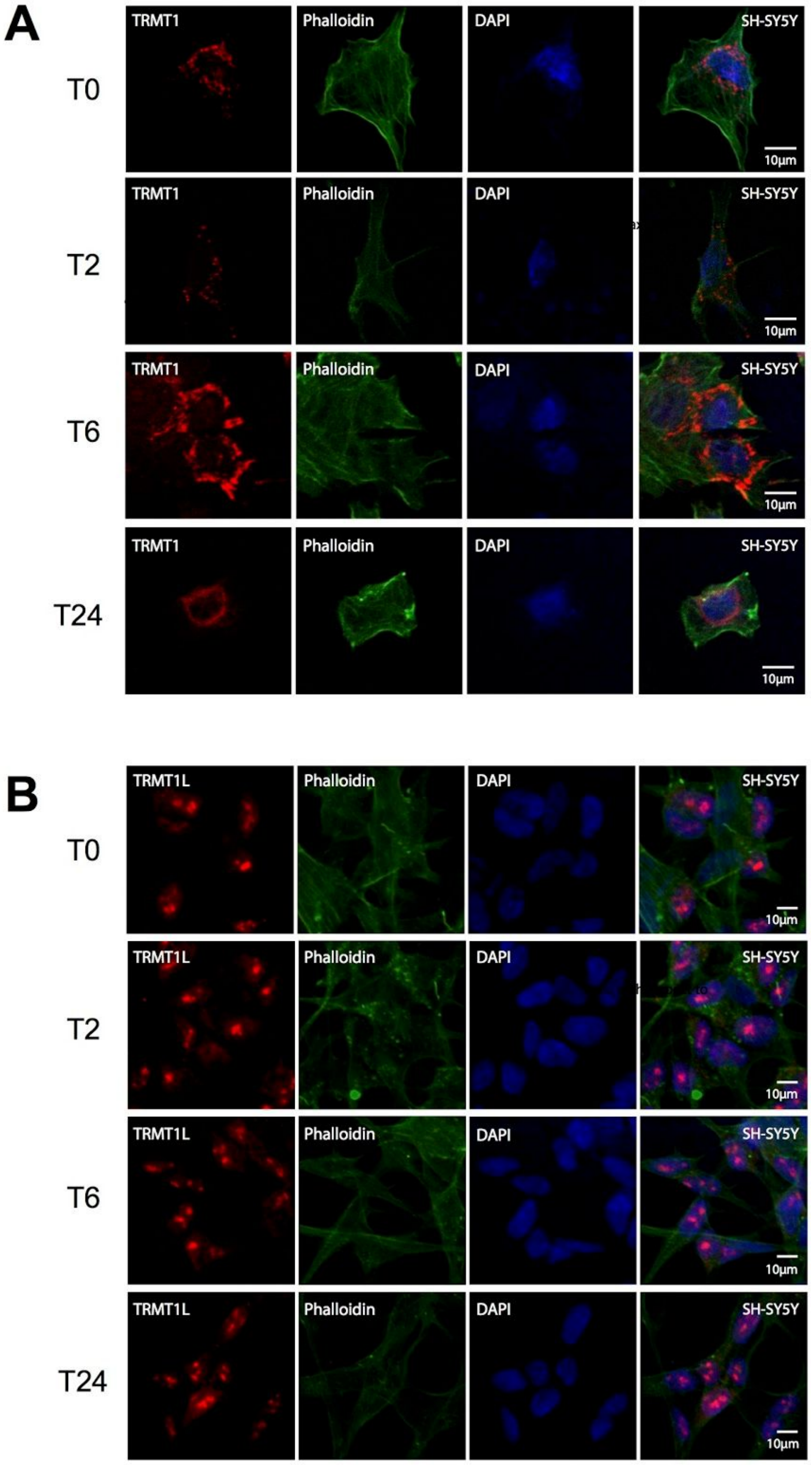

**Figure S8.** Pedigree chart of the patient-derived samples used in this study.

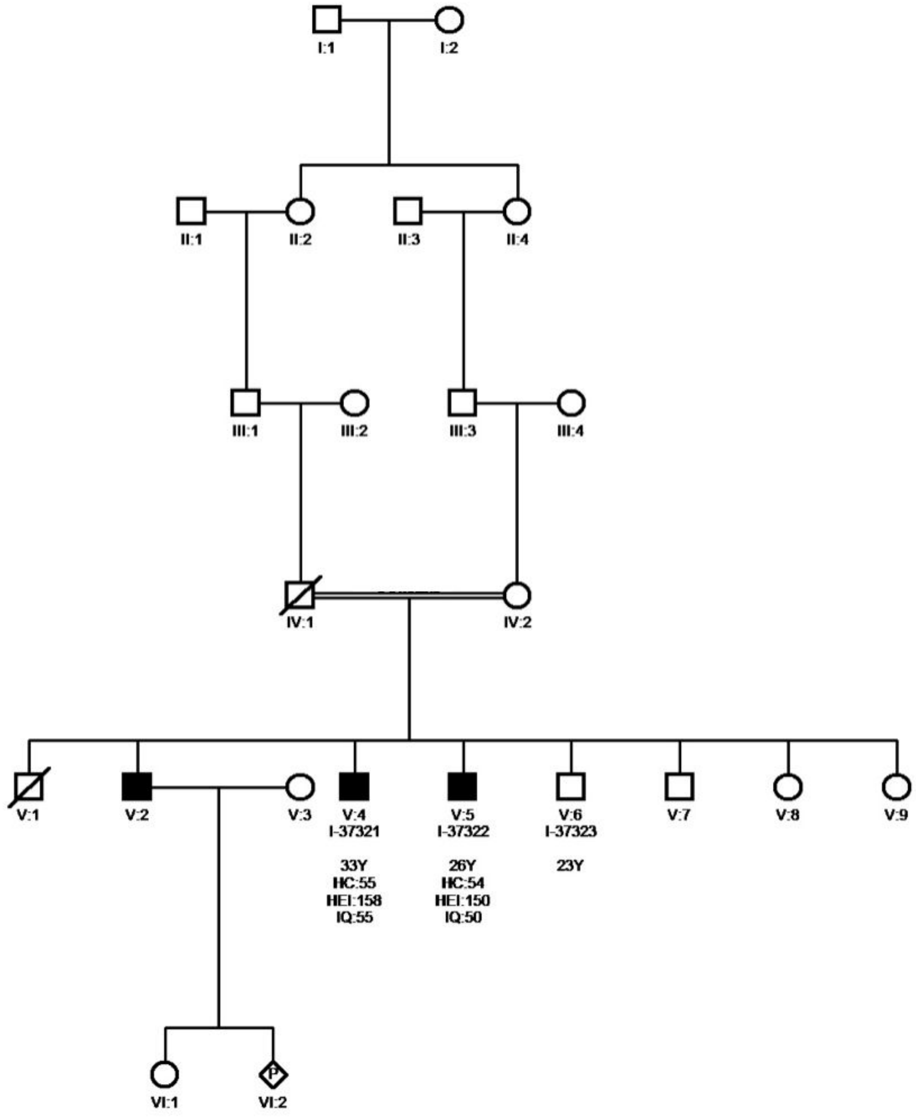

**SUPPLEMENTARY TABLES**

**Table S1. Clinical data of the patients whose blood was used to derive the LCLs which were used in this study**

| Patient ID | I-37320 | I-37321 | I-37322 |
|---|---|---|---|
| **Mutation Variant [a]** | c.1332_1333delGT; p.Y445fs | | |
| **Development** | Delay | | |
| **Age at examination (yr)** | 37 | 33 | 26 |
| **Sex** | M | M | M |
| **BW (g)** | LBW | LBW | LBW |
| **HC (cm)** | 54 | 55 | 54 |
| **Height (cm)** | 159 | 158 | 150 |
| **Weight(kg)** | 51 | 54 | 65 |
| **IQ** | 70 | 55 | 50 |
| **ID severity** | Moderate | | |
| **Speech** | Delay | | |
| **Facial dysmorphism** | Synophrys, Broad nasal bridge, hypoplastic maxilla; Prominent glabella | | |
| **Body shape** | Thin leg | | |

[a] The nomenclature is based on HGVS. All LCL cell lines used in this study were taken from a family with a pY445fs mutation (M9000114). 'c' stands for coding sequence; 'p' stands for protein sequence.

**Table S2. Oligonucleotide sequences used in this work for RT-PCR.**

| Targeted mRNA | Species | Direction | Fasta Sequence |
|---|---|---|---|
| ARC | Human | Forward | AGCGGGACCTGTACCAGAC |
| ARC | Human | Reverse | GCAGGAAACGCTTGAGCTTG |
| cFOS | Human | Forward | GGGGCAAGGTGGAACAGTTAT |
| cFOS | Human | Reverse | CCGCTTGGAGTGTATCAGTCA |
| EGR1 | Human | Forward | GCCTGCGACATCTGTGGAA |
| EGR1 | Human | Reverse | CCGCAAGTGGATCTTGGTATG |
| PGKI | Human | Forward | TACCTGCTGGCTGGATGG |
| PGKI | Human | Reverse | CCATTCCACACAATCTGCTTAG |
